# Distinct defense strategies allow different grassland species to cope with root herbivore attack

**DOI:** 10.1101/569533

**Authors:** Maxime R Hervé, Matthias Erb

## Abstract

1. Root-feeding insect herbivores are of substantial evolutionary, ecological and economical importance. Plants can resist insect herbivores through a variety of tolerance and resistance strategies. To date, few studies have systematically assessed the prevalence and importance of these strategies for root-herbivore interactions across different plant species.
2. Here, we characterize the defense strategies used by three different grassland species to cope with a generalist root herbivore, the larvae of the European cockchafer *Melolontha melolontha*.
3. Our results reveal that the different plant species rely on distinct sets of defense strategies. The spotted knapweed (*Centaurea stoebe*) resists attack by dissuading the larvae through the release of repellent chemicals. White clover (*Trifolium repens*) does not repel the herbivore, but reduces feeding, most likely through structural defenses and low nutritional quality. Finally, the common dandelion (*Taraxacum officinale*) allows *M. melolontha* to feed abundantly but compensates for tissue loss through induced regrowth.
4. Synthesis: Three co-occurring plant species have evolved different solutions to defend themselves against attack by a generalist root herbivore. The different root defense strategies may reflect distinct defense syndromes.

## Introduction

Belowground, root-feeding herbivore insects have long been known for their importance in structuring agroecosystems (Hunter, 2001). More recently, their effects on host plant interactions with aboveground insects (Biere & Goverse, 2016; Papadopoulou & van Dam, 2017), on host plant defense evolution (van Dam, 2009) and plant communities (Van der Putten, 2003) were unraveled. Given the prevalence and importance of root herbivores, an important question is how plants cope with root herbivore attack (Erb, Glauser, & Robert, 2012; Rasmann & Agrawal, 2008).

Direct plant defense strategies against root herbivores encompass resistance and tolerance (Johnson, Erb, & Hartley, 2016). Resistance can be achieved by exuding soluble or volatile repellent chemicals in the rhizosphere, and/or by producing deterrent or toxic compounds at the surface or internally (Erb et al., 2013). It can also rely on structural traits that act as deterrents or digestibility reducers (Hanley, Lamont, Fairbanks, & Rafferty, 2007). Tolerance to root herbivory has mostly been associated with the ability for compensatory growth that is accompanied by a reconfiguration of plant metabolism (Johnson, Erb, et al., 2016). Finally, indirect defense strategies work through plant-mediated reinforcement of top-down control of herbivores by the third trophic level (Turlings & Erb, 2018). Over the last years, mechanistic studies have provided detailed examples of these different traits in root-herbivore interactions (Erb et al., 2015; Johnson, Hallett, Gillespie, & Halpin, 2010; Lu et al., 2015; Rasmann et al., 2005; Robert et al., 2014). Several studies also compared defenses of different plant species against root-herbivore insects, mostly focusing on chemical resistance traits (e.g. Rasmann & Agrawal, 2011; Tsunoda, Krosse, & van Dam, 2017). However, we currently lack systematic, integrated studies that compare different direct defense traits in root-herbivore interactions across different plant species. Assessing the relative importance of different types of defenses and their combination within individual plant species into so-called plant defense-syndromes (Agrawal & Fishbein, 2006) is an important next step towards a better understanding of the ecology and evolution of root-herbivore interactions.

In the present study, we combine different experimental approaches to understand the root-defense strategies of three different, co-occurring European grassland species: the common dandelion *Taraxacum officinale* agg. (Asteraceae), the spotted knapweed *Centaurea stoebe* (Asteraceae) and white clover *Trifolium repens* (Fabaceae). All three species co-occur with a generalist root herbivore, the larva of the European cockchafer *Melolontha melolontha* (Coleoptera: Scarabeidae). *Melolontha melolontha* is native to Europe and occurs abundantly in grasslands. Its larvae develop best on this species (Hauss, 1975; Hauss & Schütte, 1976). The reasons for this preference and host suitability are unknown. Recently, it was shown that *C. stoebe* is a bad host for *M. melolontha* larvae (Huang, Zwimpfer, Hervé, Bont, & Erb, 2018). The host suitability of *T. repens* is less clear (Huang et al., 2018; Sukovata, Jaworski, Karolewski, & Kolk, 2015). Regarding potential defense strategies of the three species against root-herbivores, mechanistic work so far has mostly focused on *T. officinale*. Upon damage, *T. officinale* releases a bitter latex sap containing high amount of the sesquiterpene lactone taraxinic acid β-D-glucopyranosyl ester (TA-G) (Huber et al., 2015). High TA-G levels are associated with reduced *M. melolontha* damage, and silencing TA-G production makes *T. officinale* more attractive to *M. melolontha*, suggesting that it acts as a direct defense that deters *M. melolontha* (Bont et al., 2017; Huber et al., 2016). However, even genotypes producing high levels of TA-G are regularly attacked by *M. melolontha*, suggesting overall low resistance potential against this herbivore. Recent evidence showed that prolonged herbivory by *M. melolontha* larvae increases seed dispersal of *T. officinale*, which suggests that escaping herbivory is also part of the defense strategy of this plant species (Bont et al., 2019).

Our approach involved a set of manipulative experiments to estimate root damage and consumption by *M. melolontha* attacking the different species, root regrowth and shoot growth as tolerance mechanisms and volatile- and non-volatile attractiveness of the roots as direct resistance mechanisms. We also assessed primary metabolite levels, as well as chemical and structural defense mechanisms in the different species to determine whether low food quality may be responsible for the observed differences in resistance. By combining these measurements, we demonstrate that the three different species employ different sets of defense mechanisms to reduce or tolerate *M. melolontha* damage.

## Materials and Methods

### Plants and experimental conditions

Seeds of *C. stoebe* and *T. repens* were purchased from UFA-SAMEN (Bern, Switzerland) and Samen & Saatgut Shop (Zurich, Switzerland), respectively. For *T. officinale*, the genotype A34 was propagated in the laboratory and used for experiments. All seeds were germinated on seedling substrate and transplanted into 9 × 9 × 10 cm (L × l × H) pots filled with a mixed potting soil (‘Landerde’:peat:sand 5:4:1) after 2.5 weeks. Seedlings were transplanted individually except for *T. repens* where two seedlings were transplanted per pot to provide a sufficient amount of root material for *M. melolontha* larvae (hereafter, each pot is treated as a single replicate). Plants were used for experiments at 10 weeks after sowing. Cultivation and experiments took place in the same controlled conditions in climatic chambers: photoperiod 16:8 (light:dark), light intensity approx. 350 μmol.m^-2^.s^-1^ (supplied by Radium Bonalux NL39W 830/840 lamps), temperature 22:18 °C (day:night) and humidity 65%.

### Insects

*M. melolontha* larvae were collected from meadows in different areas of Switzerland (Table 1). Larvae were reared in controlled conditions (10 °C, darkness) in individual soil-filled plastic cups with carrot slices as food source. Second-instar (L2) and third-instar (L3) larvae were starved for five and seven days before experiments, respectively.

**Table 1.** Populations of *Melolontha melolontha* larvae used in this study. L2: second instar, L3: third instar.

| Location | Coordinates | Date of collection | Instar at collection | Instar at experiment |
| --- | --- | --- | --- | --- |
| Erstfeld | 46.82°N, 8.64°E | September 2015 | L2 | L2 |
| Kesswil | 47.60°N, 9.30°E | September 2015 | L2 | L2 |
| Bristen | 46.77°N, 8.69°E | May 2016 | L2 | L2 |
| Urmein 1 | 46.69°N, 9.41°E | May 2015 | L2 | L3 |
| Urmein 2 | 46.69°N, 9.41°E | September 2015 | L3 | L3 |
| Valzeina | 46.96°N, 9.61°E | September 2015 | L3 | L3 |

### Host suitability and estimation of root consumption

To establish the pattern of host suitability, pre-weighed *M. melolontha* larvae were individually placed with one plant for a fixed number of days. Larvae were added to plant pots into a 1cm hole near the center of the pot, and then covered with soil. At the end of the experiment, larvae were sampled back from the pots and weighed. Host suitability was assessed through larval performance, which was defined as a relative weight gain: (weight post-experiment – weight pre-experiment) / weight pre-experiment. To test for the robustness of the pattern, the experiment was conducted with two populations of L2 larvae (Erstfeld and Kesswil) and two populations of L3 larvae (Urmein 2 and Valzeina). Experiment duration was 14 days for L2 larvae, 10 days for L3 larvae. Eleven to twelve replicates were performed per population, except for Erstfeld where five to six replicates were performed due to a lower number of available larvae. To estimate root consumption, the whole root system of each plant was harvested at the end of the experiment. Soil was removed by gentle washing with tap water. Roots were then dried for 5 days at 65 °C and weighed. As a control, twelve other plants of each species were included in the experimental design. These plants were grown and harvested in the exact same conditions as the first ones but no larva was added.

### Estimation of root consumption and capacity for compensatory growth

Since root consumption estimation from the first experiment could be biased by compensatory regrowth a second experiment was conducted. Plants were grown in two stacked pots filled with the same soil. The bottom of the upper pot (‘systemic compartment’) was replaced with a fine mesh (Windhager, Switzerland). The mesh allowed roots to grow through, but restricted the herbivore larvae to the lower pot (‘attacked compartment’). Three treatments were conducted for each plant species: ‘control’, ‘larva’ (one L3 larva of population Urmein 1 placed in the attacked belowground compartment) and ‘root removal’ (mechanical removal of all roots of the attacked belowground compartment by cutting them with scissors just below the mesh, one day after the beginning of the experiment). The ‘root removal’ treatment was included to test whether plants are able to compensate for root loss without the confounding factor of different larval feeding patterns across the three species. Ten days after the beginning of the experiment, roots of each belowground compartment as well as aboveground organs were harvested separately, dried as explained above and weighed. No root could be harvested from the attacked belowground compartment of the ‘root removal’ treatment. Before harvesting of the attacked belowground compartment of the ‘larva’ treatment, damage to roots was visually assessed using a three-level damage scale: no damage except for a small spherical area around the larva (‘+’), one or several tunnels but ≤ 50% of roots removed (‘++’) or > 50% of roots removed (‘+++’). Six to seven replicates were performed per species and treatment.

### Contribution of distance and contact cues to plant resistance

Two experiments were conducted to assess whether the capacity of *C. stoebe* and *T. repens* to inhibit *M. melolontha* feeding was due to the release of repellent volatiles and/or exudates or due to contact-dependent defenses. At the beginning of the first experiment, the bottom of the pots were removed and replaced with a fine mesh (Windhager, Switzerland), then the pots were placed in a second pot filled with the same soil. The mesh was used to prevent roots from growing through and larvae from attaining the plants, while allowing exudates and volatiles to pass into the lower pot. A round piece of artificial diet (4 cm diameter, 1 cm height, 12 g, composition modified from Allsopp (1994)) was added to the lower belowground compartment, just below the mesh, and one L2 larva was placed at the bottom of the lower belowground compartment (Figure S1). After 14 days, the piece of artificial diet was recovered from the soil and damage was visually assessed using a five-level damage scale: no consumption (‘0’), 1-30% piece consumed (‘+’), 31-60% piece consumed (‘++’), 61-90% piece consumed (‘+++’), 91-100% piece consumed (‘++++’). Twelve replicates were performed per plant species (half with larvae from population Kesswil and half with larvae from population Erstfeld).

At the beginning of the second experiment, the bottom of the pots were removed and replaced with a fine mesh as in the first experiment. Root chemicals were allowed to diffuse into the lower pot over four days. At this time, one side of the lower pot was opened and this pot was fixed to another pot containing fresh soil of the same composition and moisture. A pot filled with soil was placed on the top of this second lower pot to equalize pressure in the two lower pots. At the same time, one L2 larva (population Bristen) was placed at the bottom of the pot below the plant (Figure S1). Twenty-four hours later, larvae were sampled back to assess whether they escaped form the pot containing root chemicals to the pot with fresh soil. Nineteen to twenty replicates were performed per plant species.

### Importance of root exudates for C. stoebe resistance

Since previous experiments showed chemicals released by *C. stoebe* reduce *M. melolontha* diet consumption, an additional experiment was performed to test whether this effect could be reproduced by using soluble root exudates. Exudates of *C. stoebe* and *T. officinale* were collected by placing the root system of a single intact plant (which was previously shaken gently to remove most of the surrounding soil) into 50 ml of deionized water for 3 h. The water was then centrifuged for 10 min at 3500 rpm at room temperature and the supernatant collected and freeze-dried. Four plants were used per species, which exudates were mixed after freeze-drying and re-diluted into 70 ml of deionized water. This solution was used to prepare diet pieces by mixing it with agar (size, weight and proportion of agar similar to artificial diet pieces). Pieces were then offered to single L2 larva (population Bristen) in pots filled with the same soil as in the other experiments. After seven days, the pieces were recovered from the soil and damage was visually assessed using the five-level damage scale explained above. Eight replicates were performed per species.

### Contribution of structural factors and exuded or non-exuded deterrent compounds to T. repens resistance

Since previous experiments showed that *T. repens* had a negative effect on *M. melolontha* larvae upon direct contact, but that this effect was not associated with a repellent effect of released chemicals, a series of experiments were performed on *T. repens* and *T. officinale* to test whether this effect was due to structural factors, exuded deterrent chemicals or non-exuded deterrent chemicals.

#### Structural factors

The effect of structural factors was tested with a setup based on feeding piece. Agar pieces were spiked with either 100 mg of fresh root pieces (~2 cm long) or 100 mg of fresh root powder obtained after grinding roots in liquid nitrogen. We hypothesized that grinding the roots would destroy plant structural features, including lignified cell walls, and would thus result in a food matrix in which root toughness could no longer be assessed by the larvae and thus influence their feeding behavior. Seven to twelve replicates per experiment and plant species were carried out, all of them with L2 larvae from population Bristen. To obtain a complementary chemical measure of root toughness, total lignin was quantified in roots of *T. officinale* and *T. repens*. Measurements were performed on six randomly chosen control plants per species. Lignin was extracted and quantified as described in Maia et al. (2012) based on 20 mg of dried powder.

#### Soluble exuded chemicals

Soluble exuded compounds were tested as described in the experiment comparing *T. officinale* and *C. stoebe* root exudates. The same *T. officinale* feeding pieces were used for comparison with *C. stoebe* and *T. repens*, all three plants having been tested simultaneously.

#### Soluble non-exuded chemicals

The potential of internal root-derived soluble chemicals to reduce *M. melolontha* feeding on *T. repens* was further tested by spiking agar pieces with root extracts from *T. officinale* or *T. repens*. Three kinds of extracts were prepared to test for a broad range of compound polarity: water, methanol and hexane. The water extract was prepared by continuous shaking of 1200 mg of fresh root powder (quantity equivalent to 100 mg per final feeding piece) into 40 ml of deionized water for 1 h. The extract was then centrifuged for 10 min at 3500 rpm at room temperature and the supernatant collected, then the volume completed to 70 ml using deionized water. The methanol extract was prepared by continuous shaking of 1200 mg of fresh root powder into 40 ml of methanol for 1 h. The extract was then centrifuged as above and the supernatant collected, then evaporated in a rotary vacuum evaporator at 45 °C until a volume of 5 ml was obtained. This was added to 65 ml of deionized water prior to the preparation of feeding pieces. Finally, the hexane extract was prepared by continuous shaking of 1200 mg of fresh root powder into 40 ml of hexane for 1 h. The extract was then centrifuged as above and the supernatant collected, then completely evaporated in a rotary vacuum evaporator at 45 °C. The dry residue was diluted into 5 ml of hexane:isopropanol 50:50 to improve mixing with 65 ml of deionized water during feeding piece preparation.

### Profiling of root primary metabolites

Metabolic profiling of root primary metabolites and elements was performed (i) to assess the relative nutritional quality of the different plant species, and (ii) to test whether infestation by *M. melolontha* reconfigures primary metabolism, potentially as a part of induced tolerance through resource reallocation. We assessed concentrations of essential amino acids (arginine, histidine, isoleucine, leucine, lysine, phenylalanine, threonine, valine), major simple sugars (fructose, glucose, sucrose), phytosterols (campesterol, stigmasterol, β-sitosterol) and elements (Ca, K, Mg, Na, P). Dried roots from plants of the experiment on host suitability were used as material. Measurements were performed on the same six control plants per species that were used for lignin quantification and on the twelve plants per species placed with L3 larvae from population Valzeina. Extraction and quantification of amino acids, sugars and elements was performed as described in Hervé, Delourme, Leclair, Marnet, & Cortesero (2014), Machado et al. (2013) and Neba, Newbery, & Chuyong (2016), respectively (based on 10, 10 and 30 mg of dried powder, respectively). Phytosterols were extracted according to Feng, Liu, Luo, & Tang (2015) based on 10 mg of dried powder and quantified by ultraperformance convergence chromatography – mass spectrometry. Chromatography was performed on a Waters Acquity UPC^2^ with a BEH 100 mm × 3.0 mm × 1.7 μm column, with the following parameters: column temperature 40 °C, solvent A supercritical CO2, solvent B methanol, column flow 2 ml.min^-1^, make-up solvent methanol, make-up flow 0.2 ml.min^-1^, CO2 back-pressure 2000 psi. The gradient of solvents was 0-1 min 98% A, 1-2 min linear decrease to 65% A, 2-2.5 min 65% A, 2.5-2.6 min linear increase to 98% A, 2.6-3 min equilibration at 98% A. Compounds were quantified on a Xevo G2-XS QTof high-resolution mass spectrometer with the following parameters: positive-mode ESCi multi-mode ionization (high-speed switching between electrospray ionization and atmospheric pressure chemical ionization), source temperature 120 °C, capillary voltage 3 kV, corona current 15 μA, dry gas (nitrogen) temperature 400 °C. Compounds were identified and quantified based on the following [M+H]^+^ fragments (amu): campesterol 383.3677, β-sitosterol 397.3833 and stigmasterol 395.3673. All compounds were quantified using calibration curves from pure standards.

### Data analysis

All statistical analyses were performed with the R software v. 3.4.0 (R Core Team, 2017). Pairwise comparisons of Estimated Marginal Means (EMMeans) were systematically performed if not otherwise stated, using the ‘emmeans’ package (Lenth, 2018). P-values of pairwise comparisons were always adjusted using the False Discovery Rate correction. The performance of larvae was analyzed using an ANOVA (one model per larval instar) taking into account the plant species, the larval population and the interaction between these two factors. Root consumption data were analyzed separately for each plant species using ANOVAs, which were performed separately for each larval instar in the first experiment and for each compartment (aboveground, upper belowground, lower belowground) in the second experiment. The proportion of larvae that escaped in the ‘escape experiment’ was compared between the three plant species using a likelihood ratio test applied on a generalized linear model (family: binomial, link: logit). Damage data obtained on feeding pieces or artificial diet pieces were analyzed using likelihood ratio tests applied on Cumulative Link Models (CLM), which were built using the ‘ordinal’ package (Christensen, 2018). Due to impossibility to adjust a proper CLM, root damage data were analyzed using a Kruskal-Wallis test followed by pairwise Mann-Whitney-Wilcoxon tests. Since CLMs work on latent variables which values do not make direct biological sense, medians and associated 95 % confidence intervals are systematically used for graphical representation of damage data. Primary metabolites and elements were compared between plant species using both a multivariate approach (redundancy analysis (RDA) on centered and scaled data, and associated permutation test with 9999 permutations, ‘vegan’ package (Oksanen et al., 2018)) and a univariate approach (Welch t-test for each compound, all p-values being further adjusted using a FDR correction). The same process was used to compare control and infested plants, separately for each species. Lignin content was also compared between plant species using a Welch *t*-test.

## Results

### M. melolontha larvae perform better on T. officinale than on C. stoebe and T. repens

Larval performance differed significantly between the three plant species for both L2 larvae (F_2,46_ = 9.135, *p* < 0.001) and L3 larvae (F_2,66_ = 55.542, *p* < 0.001). Overall, the L3 population Valzeina performed systematically better than the L3 population Urmein (F_1,66_ = 10.563, *p* = 0.002). No differences between the two L2 populations were observed (F_1,46_ = 0.002, *p* = 0.969). The population origin had no effect on performance patterns (L2: F_2,46_ = 0.889, *p* = 0.418, L3: F_2,66_ = 2.409, *p* = 0.098). In all cases, larval performance was better on *T. officinale* than on the two other plant species (Figure 1). Strikingly, L3 larvae did not gain any weight when feeding on *T. repens* or *C. stoebe*, suggesting the presence of strong resistance traits in these species.

**Figure 1.**
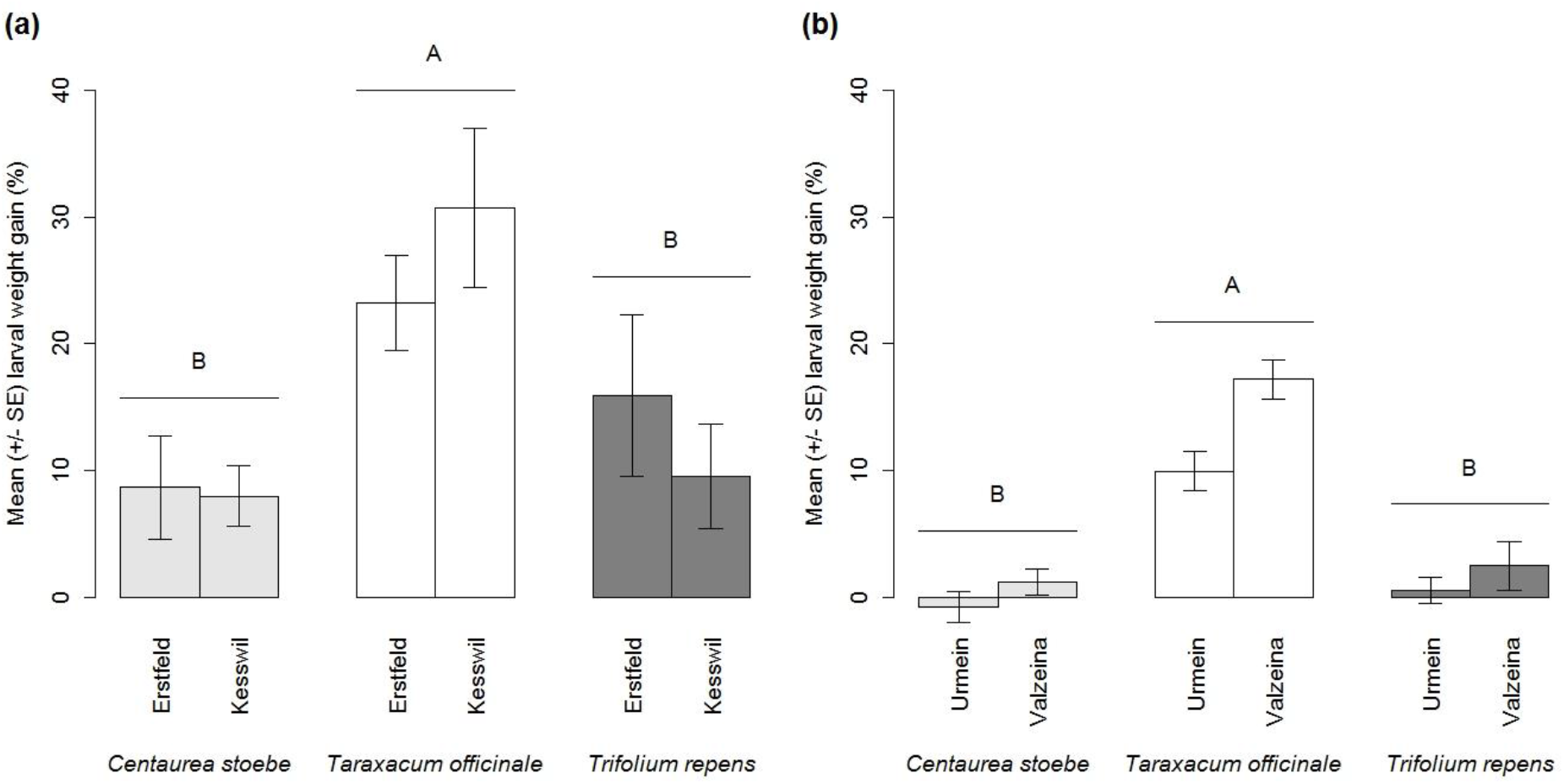
Root herbivore performance on different plant species. Performance of *Melolontha melolontha* larvae from different populations on *Centaurea stoebe, Taraxacum officinale* and *Trifolium repens*. (a) Ggrowth of second-instar larvae, (b) growth of third-instar larvae. Different letters indicate significant differences between plant species (*p* < 0.05).

### T. officinale specifically compensates for high root consumption through regrowth

No difference in *T. officinale* and *C. stoebe* root biomass was observed between control plants and plants that were infested with *M. melolontha* (*T. officinale*: L2: F_2,27_ = 0.166, *p* = 0.848, L3: F_2,33_ = 1.471, *p* = 0.244; *C. stoebe:* L2: F_2,25_ = 0.869, *p* = 0.432, L3: F_2,33_ = 0.615, *p* = 0.547) (Figure 2). By contrast, *T. repens* root dry mass was reduced significantly upon infestation by *M. melolontha* (L2: F_2,27_ = 13.494, *p* < 0.001; L3: F_2,33_ = 4.085, *p* = 0.026) (Figure 2).

**Figure 2.**
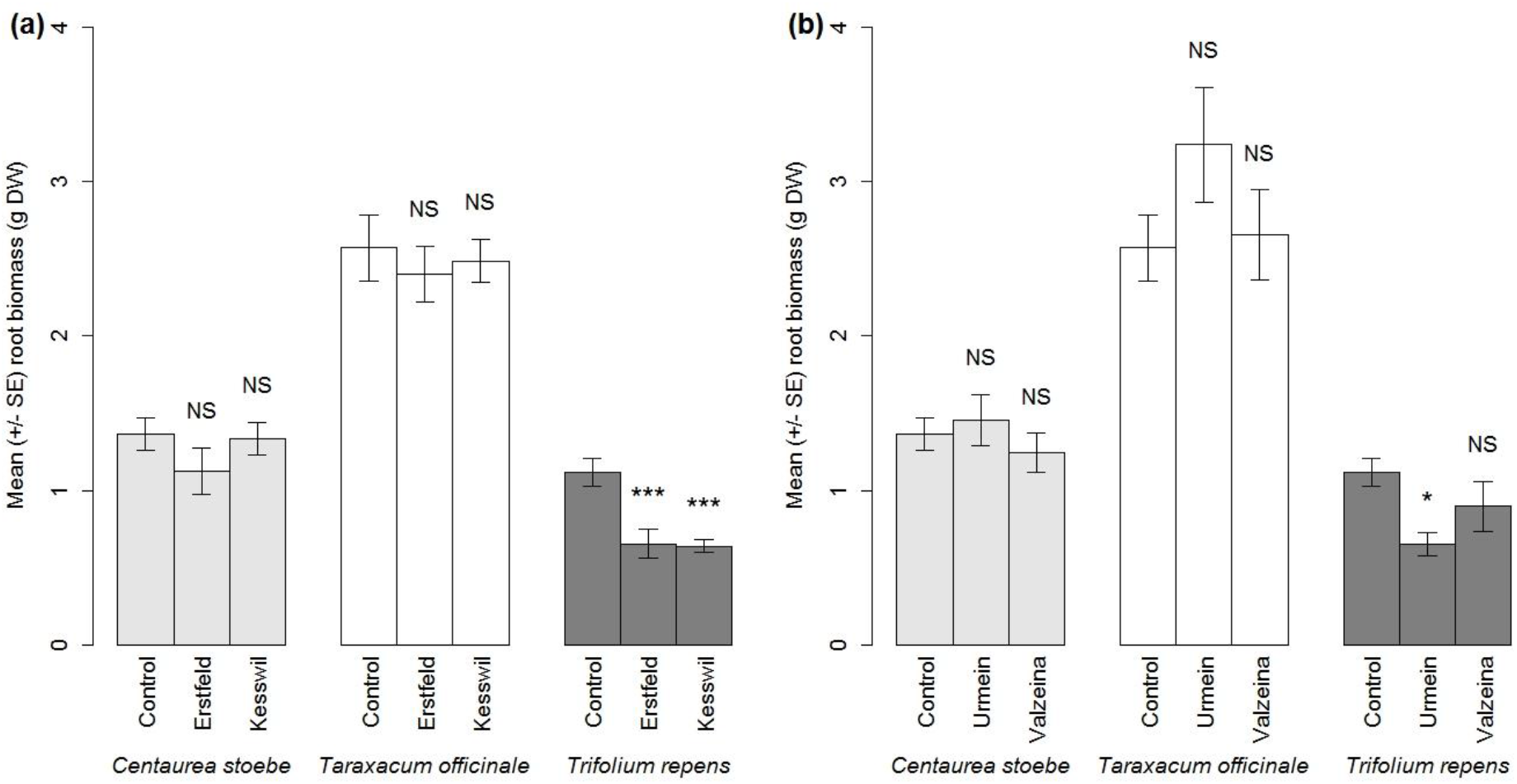
Changes in root biomass following root herbivore infestation. Root biomass of *Centaurea stoebe*, *Taraxacum officinale* and *Trifolium repens* plants that were infested with *Melolontha melolontha* larvae from different populations (Erstfeld, Kesswil, Urmein, Valzeina) or left uninfested (Control). (a) Second-instar larvae, (b) third-instar larvae. Asterisks indicate significant differences between control and infested plants (\**p* < 0.05, \*\*\**p* < 0.001). NS: not significant.

The same pattern was observed when larvae were restricted to the lower parts of the root systems of the different species. Root biomass of the attacked compartment was not different between control and infested plants for *T. officinale* (F_1,12_ = 0.887, *p* = 0.365) and *C. stoebe* (F_1,11_ = 0.000, *p* = 1.000), whereas root biomass of *T. repens* plants was significantly reduced by *M. melolontha* attack (F_1,12_ = 8.072, *p* = 0.015) (Figure 3a). Root damage scores differed between species (*χ*^2^ = 13.475, df = 2, *p* = 0.001), with *T. officinale* roots showing significantly more damage than the other two species (Figure 3b). Thus, root herbivore performance on the different species can be explained by the extent of root damage, and hence herbivore feeding, but these parameters are not reflected in final root biomass. A possible explanation for this apparent contradiction was uncovered when assessing the growth responses of the different plants upon herbivore attack and mechanical root damage. While the biomass of the shoots and the systemic roots did not change in *T. repens* in response to *M. melolontha* attack and mechanical root damage, both treatments significantly increased shoot and root biomass in *T. officinale* while in *C. stoebe* only mechanical damage increase root, but not shoot, biomass. (Figure 3c,d). Thus, *T. officinale* is most damaged and readily consumed by *M. melolontha*, but shows the strongest capacity for compensatory growth, and thus does not suffer from a reduction in vegetative growth under the given conditions. *Centaurea stoebe* on the other hand does not seem to be consumed by *M. melolontha* at all, which is reflected in the absence of root biomass increase despite capacity for compensatory growth. This plant is thus highly resistant to *M. melolontha*. Finally, *Trifolium repens* is fed upon by *M. melolontha*, as it suffers from a reduction in root biomass upon infestation, but damage remains low, suggesting that root consumption is limited. This suggests that this species is at least partially resistant to the root herbivore.

**Figure 3.**
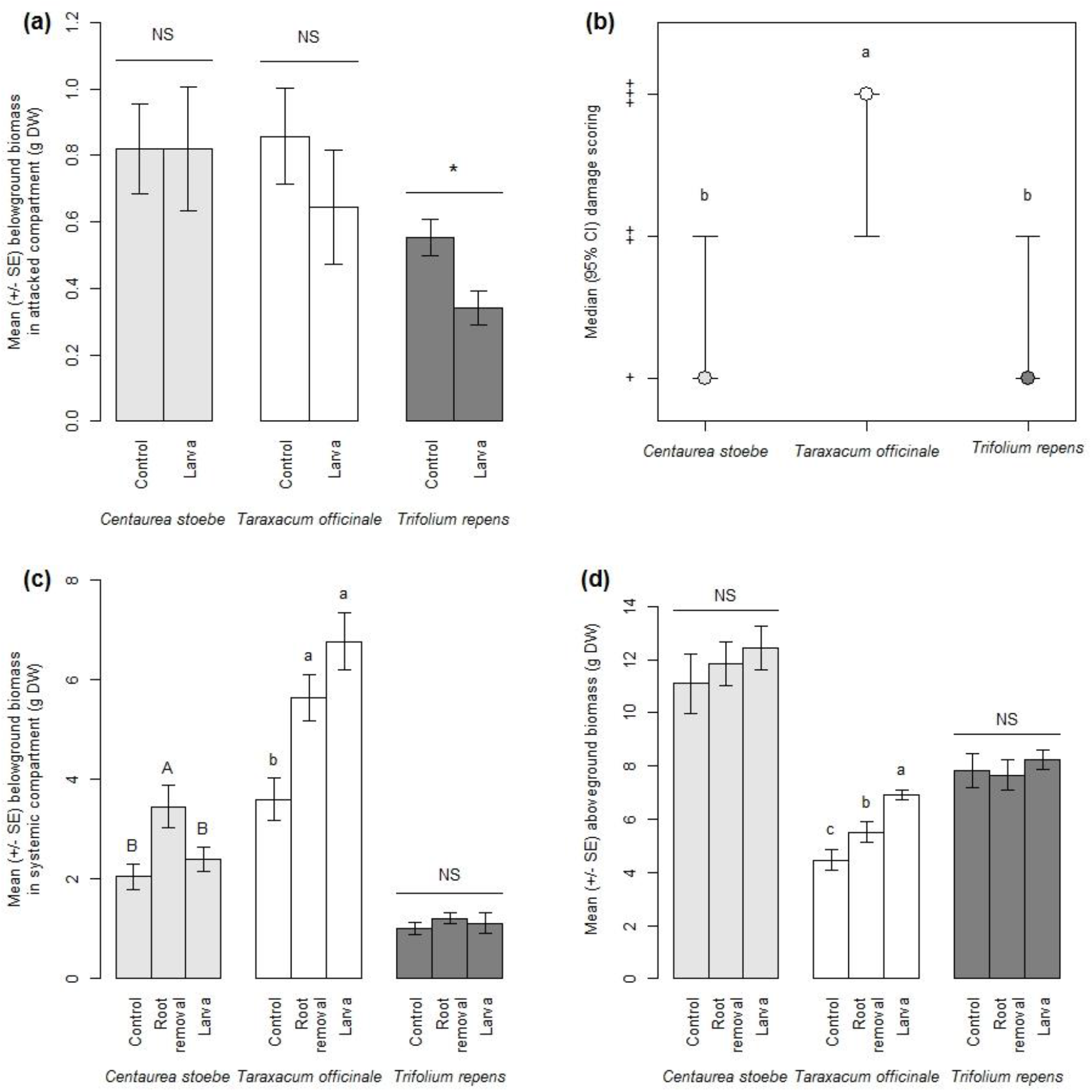
Root damage and regrowth patterns of different plant species in a split-root system. (a) Root biomass in the attacked belowground compartment in control and *Melolontha melolontha* infested plants (“Larva”). (b) Visual assessment of damage of roots within the attacked belowground compartment. Scores were ‘+’: no damage except for a small spherical area around the larva; ‘++’: one or several tunnels, but ≤ 50% of roots removed; and ‘+++’: > 50% of roots removed. (c) Root biomass in the systemic belowground compartments that were not directly attacked by *M. melolontha*. (d) Aboveground biomass. Different letters indicate significant differences between treatments or species (*p* < 0.05). Asterisks indicate significant differences between species (* *p* < 0.05).

### C. stoebe reduces M. melolontha feeding by releasing chemicals in the rhizosphere

Compared to *T. officinale*, exposure to *C. stoebe* at a distance reduced *M. melolontha* feeding on artificial diet (Figure 4a) and prompted the majority of the larvae to move away from the plant into a pot containing soil only (Figure 4b). No difference was shown between *T. officinale* and *T. repens*, either for damage (Figure 4a) or for the proportion of larvae moving away from the plant (Figure 4b). Therefore, *C. stoebe* has the capacity to repel *M. melolontha* without direct physical contact, which may contribute to its strong resistance phenotype.

**Figure 4.**
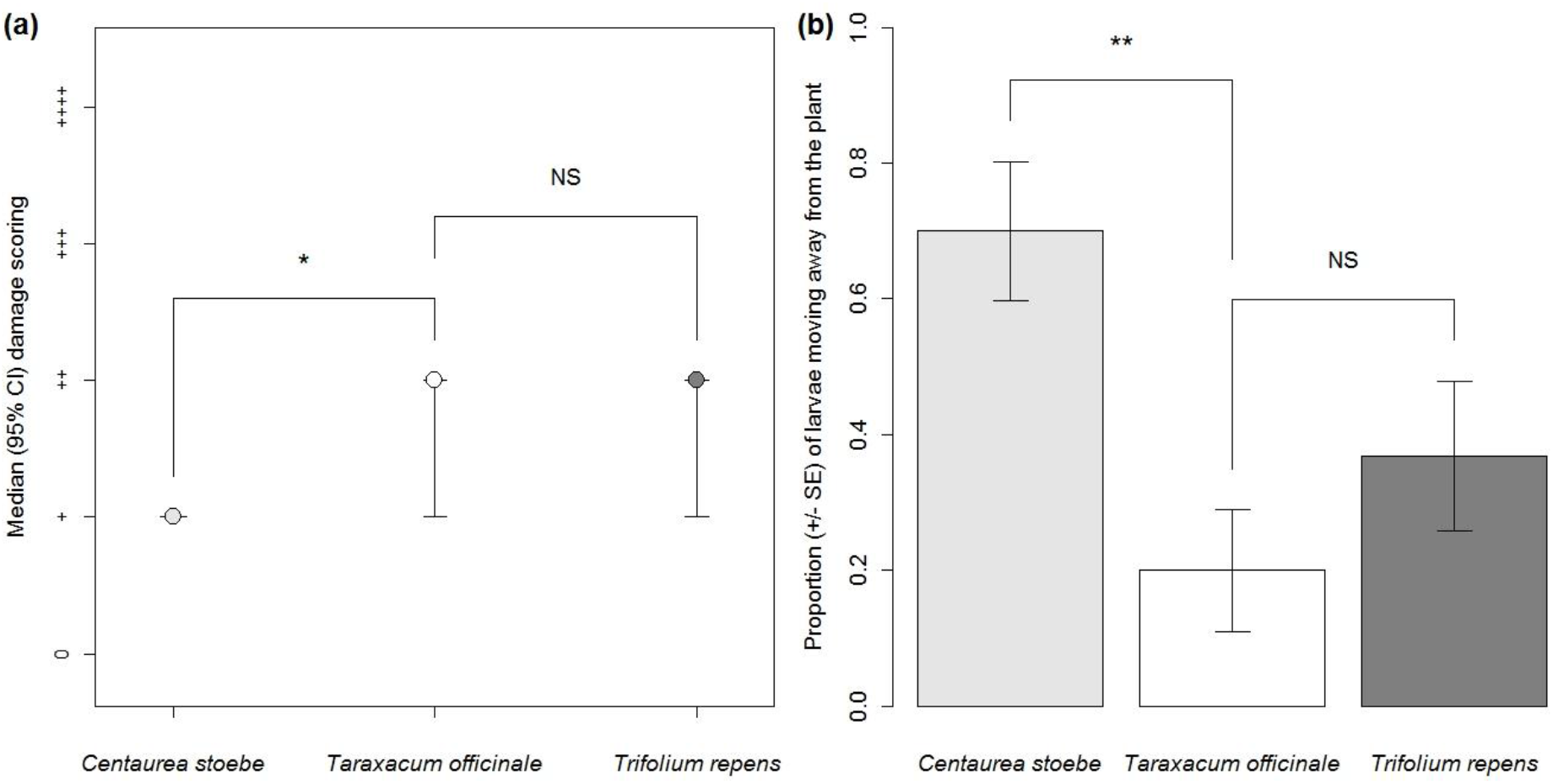
Influence of released chemicals on root-herbivore feeding behavior. (a) Feeding activity of *Melolontha melolontha* larvae on pieces of artificial diet in the vicinity of roots of the different plant species. ‘0’: no consumption; ‘+’: 1-30% piece consumed; ‘++’: 31-60% piece consumed; ‘+++’: 61-90% piece consumed; ‘++++’ 91-100% piece consumed. (b) Proportion of larvae moving away from the vicinity of the roots of the different species into a soil-filled pot without plant. Stars indicate significant differences between species (* *p* < 0.05, \*\**p* < 0.01).

### The negative effect of C. stoebe is most likely not due to soluble root exudates

No difference was observed in damage scoring of feeding pieces containing root exudates of *C. stoebe* compared to *T. officinale* (*χ*^2^ = 2.044, df = 1, *p* = 0.153). The median [95 % CI] damage scoring on *C. stoebe* was ‘+++’ [‘0’ – ‘++++’] whereas on *T. officinale* it was ‘++++’ [‘+++’ – ‘++++’].

### Structural integrity of T. repens roots is associated with lower M. melolontha root consumption

Experiments on feeding pieces showed that those containing root pieces of *T. repens* were significantly less damaged than those containing root pieces of *T. officinale*. This difference was lost when roots were ground into powder (Figure 5). Lignin content was significantly higher in roots of *T. repens* (mean ± SE 24.33 ± 1.02 μg.mg^-1^) than in *T. officinale* (18.69 ± 1.50 μg.mg^-1^) (t8.814 = −3.064, *p* = 0.014). No difference in damage was observed neither in the experiment with feeding pieces containing root exudates nor in the three experiments with feeding pieces containing root extracts (Figure 5). Thus, the higher resistance of *T. repens* is most closely associated with root structural features such as lignin-mediated toughness. Labile chemical defenses that are destroyed during root grinding and extraction may also contribute to the observed pattern.

**Figure 5.**
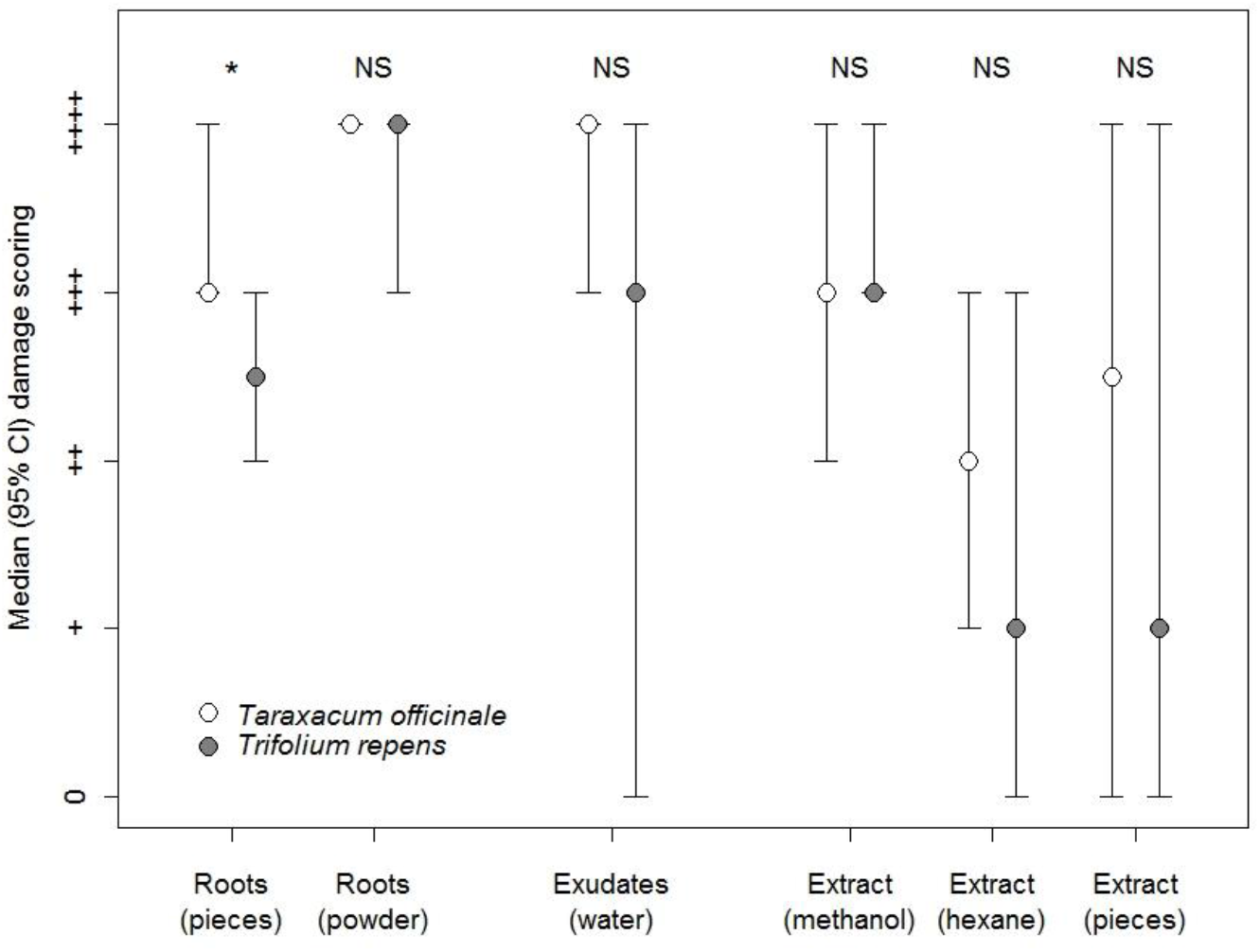
Influence of different root traits on *Melolontha melolontha* feeding. Median damage scoring of feeding pieces in a series of experiments aiming at deciphering the contribution of structural factors and phagodeterrent compounds in the negative effect of *Trifolium repens* on *Melolontha melolontha* larvae. * *p* < 0.05.

### T. repens roots are less nutritious than T. officinale roots

The RDA showed that root nutrient contents differed between *T. officinale* and *T. repens* (34.2% of constrained variance, F = 5.201, *p* = 0.006). Both multivariate and univariate approaches revealed that *T. officinale* roots contained more nutrients than *T. repens* roots (Figure 6, Table S1). The strongest differences were found for glucose (x10.9 in *Taraxacum*), fructose (x4.4), stigmatersol (x3.4) and campesterol (x2.1). There was no difference in nutrients between *T. officinale* roots and *C. stoebe* roots, both multivariately (14.4% of constrained variance, F = 1.678, *p* = 0.156) and univariately (all *p* ≥ 0.450, Table S2). Thus, the three species vary substantially in their nutrient content, with *T. officinale* roots being richer than *T. repens* roots in essential nutrients such as sugars and sterols but not different from *C. stoebe* roots.

**Figure 6.**
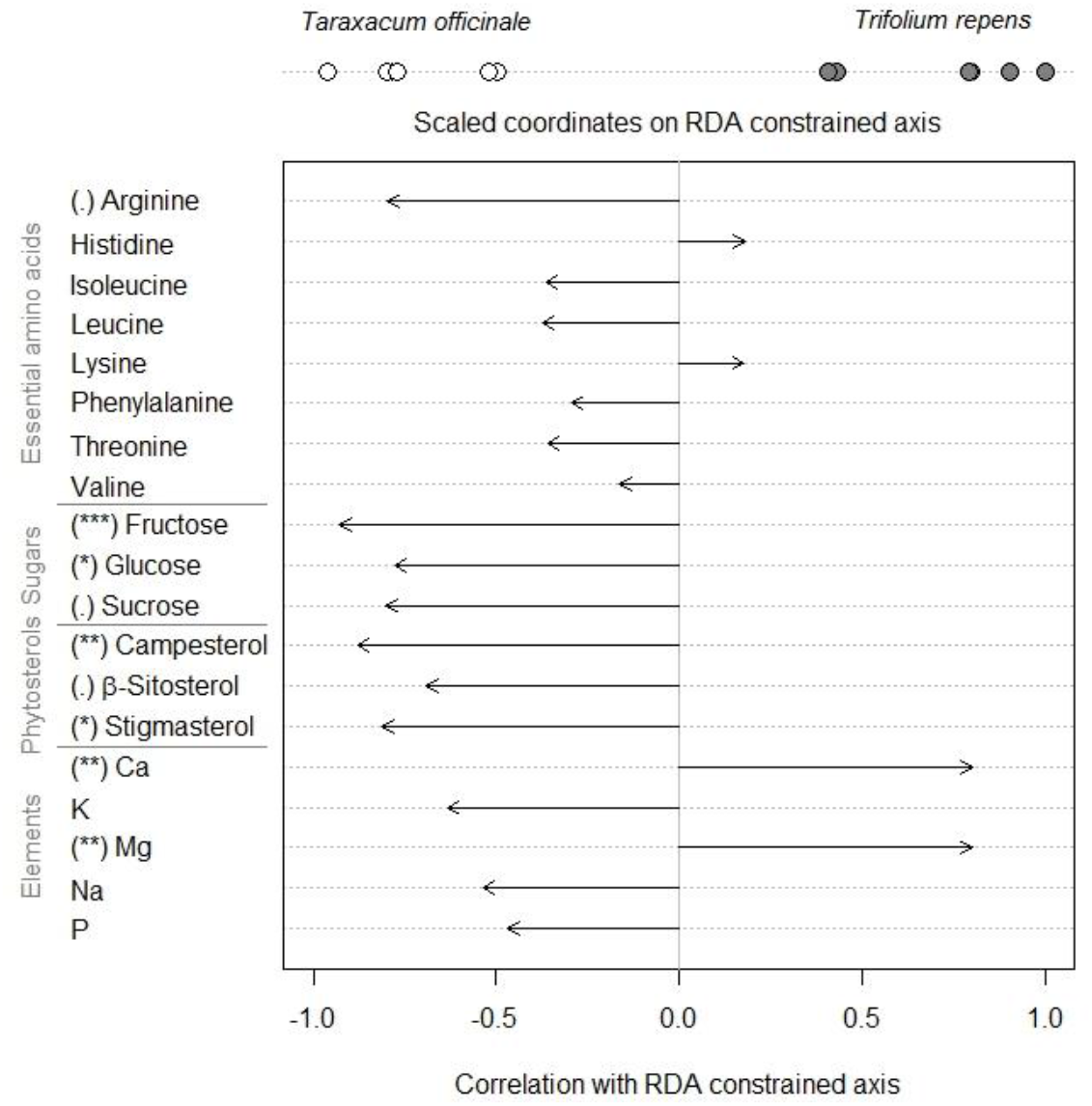
*Taraxacum officinale* roots are richer in sugars and sterols that roots of *Trifolium repens*. Redundancy analysis (RDA) performed on nutrient content of control *Taraxacum officinale* and *Trifolium repens*. Sample coordinates on the RDA constrained axis scaled to [- 1;1] and species names placed at the mean of the corresponding samples. Arrows show correlations between nutrient concentrations and the RDA constrained axis. Symbols in brackets show results of univariate tests: . *p* < 0.1, * *p* < 0.05, ** *p* < 0.01, *** *p* < 0.001. For absolute levels of nutrients, refer to Supplementary Information Table 1.

### M. melolontha attack reconfigures T. officinale primary metabolism

The RDA showed that herbivory by *M. melolontha* larvae induces significant changes in the roots’ primary metabolism of *T. officinale* (24.9% of constrained variance, F = 5.307, *p* = 0.011). The concentration of the vast majority of nutrients was lower in roots of infested plants compared to control plants (Figure 7, Table S3). The most important decrease was for simple sugars (−55.3 to −68.9%) and phytosterols (−33.4 to −46.3%). On the other hand, both multivariate and univariate approaches showed no significant change with infestation in roots of *C. stoebe* (RDA: 9.2% of constrained variance, F = 1.611, *p* = 0.142; t-tests: all *p* ≥ 0.165, Table S4) and *T. repens* (RDA: 1.5% of constrained variance, F = 0.241, *p* = 0.952; t-tests: all *p* = 0.989, Table S5).

**Figure 7.**
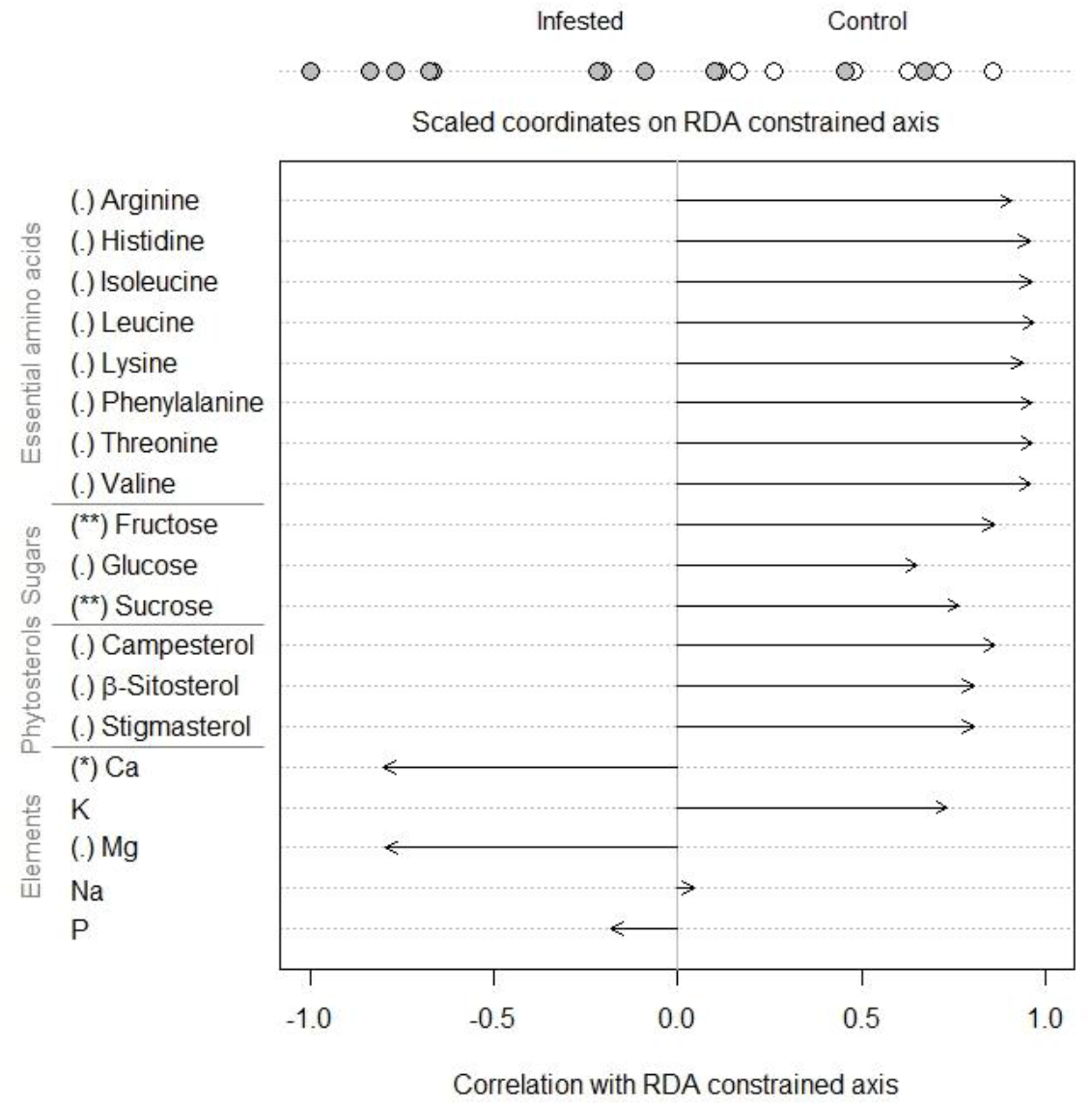
*Taraxacum officinale* roots are depleted in primary metabolites upon root herbivore attack. Redundancy analysis (RDA) performed on nutrient content of control and infested *Taraxacum officinale* plants. Sample coordinates on the RDA constrained axis scaled to [-1;1] and treatment names placed at the mean of the corresponding samples. Arrows show correlations between nutrient concentrations and the RDA constrained axis, symbols in brackets show results of univariate tests: . *p* < 0.1, * *p* < 0.05, ** *p* < 0.01. For absolute levels of nutrients, refer to Supplementary Information Table 3.

## Discussion

Plants directly defend themselves against root-feeding insects through a variety of strategies, including the storage and release of repellent chemicals, the construction of mechanical barriers and the reallocation of resources for future regrowth (Johnson, Benefer, et al., 2016; Johnson, Erb, et al., 2016). These strategies have so far mostly been investigated in isolation in individual plant species. Here, we demonstrate that three co-occurring grassland species that are threatened by the same generalist root herbivore have evolved widely different defense strategies. Below, we discuss these strategies from mechanistic and ecological points of view.

The release of repellent chemicals can be an effective strategy to avoid herbivore attack (Unsicker, Kunert, & Gershenzon, 2009). We found that, although *C. stoebe* contains high levels of nutrients similar to *T. officinale*, it does not support *M. melolontha* growth, an effect that is associated with low damage and root removal. Thus, we hypothesized that *C. stoebe* exhibits strong, almost qualitative resistance against *M. melolontha*. Indeed, M. melolontha feeding is inhibited even in the absence of direct root contact, and the larvae actively try to move away from *C. stoebe* This is one of a very few examples of repellent compounds acting at distance belowground (Johnson & Nielsen, 2012). Semi-artificial diets incorporating root exudates showed no adverse effect on *M. melolontha*, suggesting that repellent volatiles may be involved. *Melolontha melolontha* possess numerous olfactory receptors and is able to detect a diversity of volatile compounds (Eilers, Talarico, Hansson, Hilker, & Reinecke, 2012). Moreover, volatile-oriented behavior has been proven in two close relative species, *M. hippocastani* (Weissteiner et al., 2012) and *Costelytra zealandica* (Rostás, Cripps, & Silcock, 2015). The repellent volatiles of *C. stoebe* are not identified yet. However, it is known that volatile bouquets emitted by roots of *C. stoebe* are dominated by high amounts of sesquiterpenes, among a diversity of other compounds (Gfeller et al., 2019). These terpenes have so far been associated with an increase rather than a decrease of *M. melolontha* growth on neighboring plants (Huang, Gfeller, & Erb, 2019). Whether the reduction in feeding observed here is dose-dependent or due to other volatile chemical cues, and whether labile soluble exudates may play a role remains to be determined. Taken together, our profiling suggests that *C. stoebe* is protected against *M. melolontha* through the release of repellent chemicals rather than strong regrowth capacity or poor nutritional value.

Apart from the release of chemicals, plants can protect their tissues through internal structural and chemical resistance traits. We found that *T. repens* is resistant to *M. melolontha* as *C. stoebe*, but that this trait is not associated with repellency from a distance. The semi-artificial diet further showed that neither root exudates, nor soluble internal chemicals can explain this resistance. Instead, intact root pieces seem to be disliked by *M. melolontha*, a pattern that is associated with high levels of root lignin in *T. repens*. As lignin directly contributes to tissue toughness, it is conceivable that higher lignification may stop *M. melolontha* from feeding on *T. repens* (Johnson, Benefer, et al., 2016). Lignin content was documented to increase root toughness and *Agriotes* spp. resistance in tobacco (Johnson et al., 2010). Additionally, our metabolic profiling showed that the nutritional quality of *T. repens* is substantially lower than that of *T. officinale*. Thus, apart from structural defenses, low nutrient levels may contribute to the low performance of *M. melolontha* on *T. repens*. Together, these results suggest that *T. repens* becomes resistant to M. melolontha because of low digestibility associated with high lignin and low nutrient contents.

The performance of the herbivore was the best on *T. officinale*, confirming that this species is a good host for *M. melolontha* larvae (Hauss, 1975; Hauss & Schütte, 1976). This is in line with the fact that *T. officinale* roots re nutrient rich. In an interspecific study, Sukovata et al. (2015) showed that *M. melolontha* larvae grow better on plants that are more sugar-rich. Although latex defenses protect *T. officinale* to a certain degree by prompting larvae to move to congeners with lower latex defense levels (Bont et al., 2017; Huber et al., 2016), this form of resistance is not sufficient for *T. officinale* to avoid attack by *M. melolontha* in the field. Instead, as shown here, *T. officinale* has a high capacity to compensate for root loss by increasing root growth in undamaged parts of the root system as well as shoot growth. This response is associated with a substantial reduction of primary metabolites in the attacked roots, which could have been selected as a reallocation to aboveground organs favoring tolerance, a sequestration strategy to protect nutrients away from the tissues under attack and/or a direct defense strategy decreasing nutritional quality for the herbivore, as hypothesized in cases of generalist herbivores with low mobility (Berenbaum, 1995; Johnson, Erb, et al., 2016). Taken together, *T. officinale* seems to be highly nutritious and little defended towards *M. melolontha*, but seems to be able to tolerate attack through compensatory growth.

Of note, the defense strategies of the plant species tested in this study closely match the defense syndromes described for aboveground traits of milkweeds by Agrawal & Fishbein (2006). *Centaurea stoebe* seems to follow ‘Nutrition and defense’, with good nutritional quality but strong resistance traits repelling *M. melolontha* larvae. *Trifolium repens* would fit into the category ‘Low nutritional quality’, with structural defenses combined with low nutritional quality. *Taraxacum officinale* seems to follow a ‘Tolerance/escape’ strategy, with important abilities to compensate for root loss and, as shown by Bont et al. (2019), increased seed dispersal. The fact that tolerance is expected to exert no selection pressure on herbivores (Weis & Franks, 2006) may explain why *T. officinale* is the preferred host plant of *M. melolontha* and why there is a positive historical relationship between *M. melolontha* and *T. officinale* abundance in European grasslands (Schütte, 1996). Interestingly, *T. officinale* is also one of the preferred host plants of wireworms, that co-occur with *M. melolontha* in European grasslands (Wallinger et al., 2014). This suggests that the defense strategy of *T. officinale* against generalist root herbivores might be independent of the herbivore species. From the perspective of the herbivore, our work raises questions regarding the evolution of host preference in generalist root herbivores. Could it be that host preferences in these insect species are driven by intrinsic defense strategies of their hosts, resulting in preferences for tolerant over resistant plants over evolutionary time? If this were the case, we would expected generalist root herbivores to accumulate on tolerant plants in the field. The hypothesis that accumulation of generalists predicts the defense syndrome of plants within natural communities remains to be tested.

## Authors’ Contributions

MRH and ME conceived the ideas and designed methodology; MRH collected the data; MRH analyzed the data; MRH and ME interpreted the data and wrote the manuscript.

## Acknowledgements

We thank Zoe Bont, Wei Huang, Noëlle Schenk, Elias Zwimpfer, Gabriel Ulrich, Marlise Zimmermann and Julia Fricke for field collection of *M. melolontha* larvae, rearing of larvae and technical assistance during experiments, as well as the gardeners of the IPS for their help with plant cultivation. This work was supported by a CJS grant from INRA, the Swiss National Science Foundation (grants #153517 and 157884) and the University of Bern.

## Data Accessibility

The data of this manuscript has been deposited on Dryad [to be inserted at a later date].

