## Supplementary Information for "Distinct defense strategies allow different grassland species to cope with root herbivore attack"

Hervé MR & Erb M

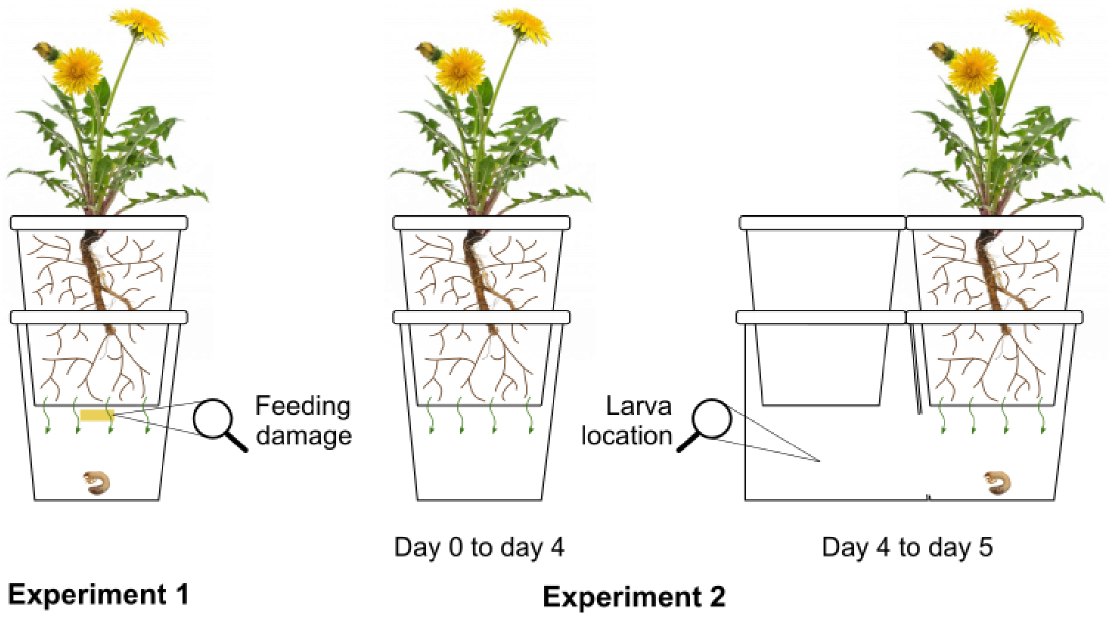


**Figure S1.** Experimental setups to test for the effect of released root chemicals on the feeding behavior of *Melolontha melolontha* larvae. In experiment 1, the impact of released root chemicals of three grassland species on diet consumption was measured. In experiment 2, the impact of released root chemicals on the probability of *M. melolontha* to leave the vicinity of the root systems of the different grassland species was quantified. Green arrows show diffusion of root chemicals. Yellow rectangle is a piece of artificial diet. A dandelion plant is shown for illustrative purposes. Experiments were carried out with *Taraxacum officinale*, *Trifolium repens* and *Centaurea stoebe*.

**Table S1.** Mean (SE) concentration of nutrients in control *Taraxacum officinale* and *Trifolium repens* plants (µg.mg$^{-1}$ DW), and results of Welch *t*-tests (FDR adjusted *p*-values).

| | *T. officinale* | *T. repens* | *t* | df | *p* |
|---|---|---|---|---|---|
| Essential amino acids | | | | | |
| Arginine | 4.82 (0.59) | 2.83 (0.47) | 2.658 | 9.521 | 0.059 |
| Histidine | 1.52 (0.12) | 1.96 (0.30) | -1.359 | 6.682 | 0.306 |
| Isoleucine | 3.96 (0.32) | 3.70 (0.51) | 0.430 | 8.436 | 0.769 |
| Leucine | 6.19 (0.44) | 5.76 (0.79) | 0.468 | 7.839 | 0.769 |
| Lysine | 3.87 (0.33) | 5.04 (0.77) | -1.396 | 6.750 | 0.306 |
| Phenylalanine | 3.69 (0.27) | 3.57 (0.50) | 0.203 | 7.792 | 0.844 |
| Threonine | 5.00 (0.35) | 4.72 (0.57) | 0.416 | 8.328 | 0.769 |
| Valine | 5.33 (0.41) | 5.52 (0.73) | -0.227 | 7.858 | 0.844 |
| Simple sugars | | | | | |
| Fructose | 14.71 (1.16) | 3.32 (0.82) | 7.999 | 9.004 | < 0.001 |
| Glucose | 16.86 (3.47) | 1.54 (0.72) | 4.321 | 5.423 | 0.020 |
| Sucrose | 38.74 (3.71) | 20.44 (5.20) | 2.865 | 9.040 | 0.050 |
| Phytosterols | | | | | |
| Campesterol | 0.05 (0.005) | 0.02 (0.003) | 4.361 | 9.474 | 0.008 |
| β-Sitosterol | 0.40 (0.06) | 0.22 (0.03) | 2.599 | 7.462 | 0.071 |
| Stigmasterol | 0.28 (0.04) | 0.08 (0.03) | 3.887 | 8.480 | 0.016 |
| Elements | | | | | |
| Ca | 19.01 (0.58) | 24.05 (0.94) | -4.586 | 8.306 | 0.008 |
| K | 13.80 (1.39) | 10.96 (1.06) | 1.628 | 9.348 | 0.236 |
| Mg | 4.18 (0.33) | 6.20 (0.20) | -5.320 | 8.208 | 0.006 |
| Na | 1.89 (0.28) | 1.24 (0.13) | 2.094 | 7.112 | 0.140 |
| P | 1.01 (0.09) | 0.87 (0.07) | 1.298 | 9.307 | 0.306 |

**Table S2.** Mean (SE) concentration of nutrients in control *Taraxacum officinale* and *Centaurea stoebe* plants ($\mu g.mg^{-1}$ DW), and results of Welch *t*-tests (FDR adjusted *p*-values).

| | *T. officinale* | *C. stoebe* | *t* | df | *p* |
|---|---|---|---|---|---|
| Essential amino acids | | | | | |
| Arginine | 4.82 (0.59) | 2.97 (0.17) | -3.024 | 5.869 | 0.450 |
| Histidine | 1.52 (0.12) | 1.45 (0.09) | -0.476 | 8.948 | 0.732 |
| Isoleucine | 3.96 (0.32) | 3.48 (0.21) | -1.228 | 8.541 | 0.487 |
| Leucine | 6.19 (0.44) | 5.51 (0.32) | -1.247 | 9.174 | 0.487 |
| Lysine | 3.87 (0.33) | 3.42 (0.20) | -1.164 | 8.146 | 0.487 |
| Phenylalanine | 3.69 (0.27) | 3.35 (0.20) | -0.991 | 9.045 | 0.508 |
| Threonine | 5.00 (0.35) | 4.38 (0.22) | -1.475 | 8.368 | 0.487 |
| Valine | 5.33 (0.41) | 4.81 (0.26) | -1.076 | 8.431 | 0.493 |
| Simple sugars | | | | | |
| Fructose | 14.71 (1.16) | 13.05 (3.32) | -0.470 | 6.207 | 0.487 |
| Glucose | 16.86 (3.47) | 10.45 (3.19) | -1.360 | 9.931 | 0.450 |
| Sucrose | 38.74 (3.71) | 29.04 (3.74) | -1.841 | 9.999 | 0.867 |
| Phytosterols | | | | | |
| Campesterol | 0.05 (0.005) | 0.05 (0.01) | 0.174 | 6.380 | |
| β-Sitosterol | 0.40 (0.06) | 0.26 (0.04) | -1.952 | 9.033 | 0.450 |
| Stigmasterol | 0.28 (0.04) | 0.30 (0.08) | 0.198 | 7.480 | 0.867 |
| Elements | | | | | |
| Ca | 19.01 (0.58) | 19.81 (0.71) | 0.878 | 9.591 | 0.545 |
| K | 13.80 (1.39) | 12.08 (1.58) | -0.818 | 9.831 | 0.548 |
| Mg | 4.18 (0.33) | 5.49 (0.58) | 1.965 | 7.853 | 0.450 |
| Na | 1.89 (0.28) | 2.79 (0.44) | 1.735 | 8.595 | 0.450 |
| P | 1.01 (0.09) | 0.86 (0.10) | -1.140 | 9.742 | 0.487 |

**Table S3.** Mean (SE) concentration of nutrients in *Taraxacum officinale* control and infested plants (µg.mg$^{-1}$ DW), and results of Welch *t*-tests (FDR adjusted *p*-values).

| | Control | Infested | *t* | df | *p* |
|---|---|---|---|---|---|
| Essential amino acids | | | | | |
| Arginine | 4.82 (0.59) | 3.13 (0.34) | 2.499 | 8.429 | 0.062 |
| Histidine | 1.52 (0.12) | 1.13 (0.11) | 2.356 | 12.433 | 0.062 |
| Isoleucine | 3.96 (0.32) | 2.92 (0.28) | 2.426 | 12.123 | 0.062 |
| Leucine | 6.19 (0.44) | 4.58 (0.45) | 2.561 | 13.927 | 0.062 |
| Lysine | 3.87 (0.33) | 3.00 (0.29) | 1.980 | 12.525 | 0.083 |
| Phenylalanine | 3.69 (0.27) | 2.75 (0.27) | 2.457 | 13.434 | 0.062 |
| Threonine | 5.00 (0.35) | 3.74 (0.33) | 2.619 | 12.869 | 0.062 |
| Valine | 5.33 (0.41) | 4.02 (0.35) | 2.446 | 11.916 | 0.062 |
| Simple sugars | | | | | |
| Fructose | 14.71 (1.16) | 6.58 (1.07) | 5.143 | 12.875 | 0.004 |
| Glucose | 16.86 (3.47) | 5.24 (1.06) | 3.201 | 5.948 | 0.062 |
| Sucrose | 38.74 (3.71) | 16.82 (2.24) | 5.059 | 8.790 | 0.007 |
| Phytosterols | | | | | |
| Campesterol | 0.05 (0.005) | 0.04 (0.01) | 2.012 | 15.998 | 0.078 |
| β-Sitosterol | 0.40 (0.06) | 0.23 (0.05) | 2.122 | 11.751 | 0.076 |
| Stigmasterol | 0.28 (0.04) | 0.15 (0.04) | 2.216 | 12.988 | 0.072 |
| Elements | | | | | |
| Ca | 19.01 (0.58) | 22.82 (1.08) | -3.128 | 15.427 | 0.043 |
| K | 13.80 (1.39) | 12.85 (0.93) | 0.568 | 9.596 | 0.583 |
| Mg | 4.18 (0.33) | 5.28 (0.40) | -2.115 | 15.512 | 0.746 |
| Na | 1.89 (0.28) | 1.68 (0.14) | 0.674 | 7.639 | 0.571 |
| P | 1.01 (0.09) | 1.08 (0.05) | -0.639 | 8.078 | 0.571 |

**Table S4.** Mean (SE) concentration of nutrients in *Centaurea stoebe* control and infested plants ($\mu g.mg^{-1}$ DW), and results of Welch *t*-tests (FDR adjusted *p*-values).

| | Control | Infested | *t* | df | *p* |
|---|---|---|---|---|---|
| Essential amino acids | | | | | |
| Arginine | 2.97 (0.17) | 3.02 (0.34) | -0.120 | 15.214 | 0.906 |
| Histidine | 1.45 (0.09) | 1.54 (0.14) | -0.536 | 15.920 | 0.759 |
| Isoleucine | 3.48 (0.21) | 3.72 (0.31) | -0.642 | 15.999 | 0.725 |
| Leucine | 5.51 (0.32) | 5.87 (0.48) | -0.635 | 15.999 | 0.725 |
| Lysine | 3.42 (0.20) | 3.93 (0.41) | -1.101 | 14.859 | 0.725 |
| Phenylalanine | 3.35 (0.20) | 3.64 (0.29) | -0.818 | 16.000 | 0.725 |
| Threonine | 4.38 (0.22) | 4.51 (0.34) | -0.317 | 15.988 | 0.844 |
| Valine | 4.81 (0.26) | 5.02 (0.40) | -0.448 | 15.981 | 0.784 |
| Simple sugars | | | | | |
| Fructose | 13.05 (3.32) | 7.34 (0.78) | 1.675 | 5.560 | 0.725 |
| Glucose | 10.45 (3.19) | 5.01 (0.98) | 1.629 | 5.952 | 0.725 |
| Sucrose | 29.04 (3.74) | 14.26 (1.58) | 3.636 | 6.848 | 0.165 |
| Phytosterols | | | | | |
| Campesterol | 0.05 (0.01) | 0.04 (0.01) | 0.998 | 6.930 | 0.725 |
| β-Sitosterol | 0.26 (0.04) | 0.22 (0.02) | 0.674 | 8.371 | 0.725 |
| Stigmasterol | 0.31 (0.08) | 0.23 (0.03) | 0.859 | 6.690 | 0.725 |
| Elements | | | | | |
| Ca | 19.81 (0.71) | 20.59 (0.89) | -0.683 | 15.566 | 0.725 |
| K | 12.08 (1.58) | 11.66 (0.54) | 0.251 | 6.220 | 0.855 |
| Mg | 5.49 (0.58) | 6.22 (0.22) | -1.171 | 6.404 | 0.725 |
| Na | 2.79 (0.44) | 2.17 (0.11) | 1.387 | 5.631 | 0.725 |
| P | 0.86 (0.10) | 0.99 (0.03) | -1.194 | 0.859 | 0.725 |

**Table S5.** Mean (SE) concentration of nutrients in *Trifolium repens* control and infested plants ($\mu g.mg^{-1}$ DW), and results of Welch *t*-tests (FDR adjusted *p*-values).

| | Control | Infested | *t* | df | *p* |
|---|---|---|---|---|---|
| Essential amino acids | | | | | |
| Arginine | 2.83 (0.47) | 2.94 (0.27) | -0.214 | 8.392 | 0.989 |
| Histidine | 1.96 (0.30) | 1.96 (0.17) | -0.015 | 8.339 | 0.989 |
| Isoleucine | 3.70 (0.51) | 3.75 (0.30) | -0.088 | 8.568 | 0.989 |
| Leucine | 5.76 (0.79) | 5.85 (0.47) | -0.097 | 8.748 | 0.989 |
| Lysine | 5.04 (0.77) | 5.14 (0.44) | -0.115 | 8.307 | 0.989 |
| Phenylalanine | 3.57 (0.50) | 3.60 (0.29) | -0.042 | 8.577 | 0.989 |
| Threonine | 4.72 (0.57) | 4.75 (0.35) | -0.052 | 8.887 | 0.989 |
| Valine | 5.52 (0.73) | 5.48 (0.43) | 0.052 | 8.514 | 0.989 |
| Simple sugars | | | | | |
| Fructose | 3.32 (0.82) | 3.50 (0.50) | -0.190 | 8.870 | 0.989 |
| Glucose | 1.54 (0.72) | 1.70 (0.60) | -0.169 | 11.936 | 0.989 |
| Sucrose | 20.44 (5.20) | 19.36 (4.50) | 0.158 | 12.193 | 0.989 |
| Phytosterols | | | | | |
| Campesterol | 0.02 (0.003) | 0.02 (0.001) | 0.938 | 6.675 | 0.989 |
| β-Sitosterol | 0.22 (0.03) | 0.19 (0.02) | 0.999 | 7.933 | 0.989 |
| Stigmasterol | 0.08 (0.03) | 0.06 (0.01) | 0.647 | 7.757 | 0.989 |
| Elements | | | | | |
| Ca | 24.05 (0.94) | 23.28 (1.55) | 0.424 | 15.848 | 0.989 |
| K | 10.96 (1.06) | 10.98 (0.55) | -0.014 | 7.863 | 0.989 |
| Mg | 6.20 (0.20) | 6.71 (0.27) | -1.507 | 15.906 | 0.989 |
| Na | 1.24 (0.13) | 1.26 (0.12) | -0.121 | 12.624 | 0.989 |
| P | 0.87 (0.07) | 0.86 (0.07) | 0.156 | 14.431 | 0.989 |
